# Altered developmental profile of the cortical head-direction signal in a rat model of Fragile X Syndrome

**DOI:** 10.1101/2025.01.09.632139

**Authors:** Noah Moore, Adrian J. Duszkiewicz, Antonis Asiminas, Paul A. Dudchenko, Adrien Peyrache, Emma R. Wood

## Abstract

Fragile X Syndrome (FXS) is a common inherited monogenic cause of autism spectrum disorder and intellectual disability (ASD/ID), but the neural mechanisms underlying its symptoms remain unclear. Here, we investigate the development of the head-direction (HD) system in a rat model of FXS (*Fmr1^−/y^* rats) and its role in spatial cognition. Using high-density silicon probes, we recorded neuronal activity in the postsubiculum (PoSub), a cortical hub of the HD system, during exploration and sleep in juvenile and adult *Fmr1^−/y^* and wild-type (WT) rats. Surprisingly, juvenile *Fmr1^−/y^* rats exhibited enhanced HD tuning characterized by sharper directional tuning curves and improved stability of the HD signal in relation to the external environment. This enhancement was intrinsic to the HD circuit and persisted across behavioral and sleep states. However, by adulthood, HD tuning in *Fmr1^−/y^* rats became unstable, with increased drift in the allocentric reference frame despite intact intrinsic HD dynamics. These findings reveal a transient enhancement of spatial coding in the HD system of juvenile *Fmr1^−/y^* rats, followed by a degradation in adulthood, mirroring the developmental regression often observed in autism. Overall, our work reveals network-level dysfunctions in FXS and their developmental trajectories.

## Introduction

Autism spectrum disorder and intellectual disability (ASD/ID) are comorbid conditions characterized by abnormalities in early cognitive development that persist into adulthood. In many cases, their symptoms are linked to *de novo* or inherited mutations in genes involved in neuronal function ^1^. With the aid of genetic rodent models, much progress has been made in uncovering how these single-gene mutations affect intracellular processes like excitability, synaptic transmission and plasticity ^2^. However, how such synaptic phenotypes give rise to cognitive symptoms in ASD/ID at a systems level is not well understood.

The neurodevelopmental nature of ASD/ID necessitates quantification of systems-level phenotypes during early postnatal development - a task aided by the recent introduction of rat ASD/ID models ^3–8^. Indeed, compensation mechanisms that emerge throughout adolescence often occlude the primary circuit dysfunctions that give rise to ASD/ID phenotypes ^9^. In recent years, the problem has been addressed by focusing on primary sensory systems ^10–12^, in which neural tuning to sensory features can be quantified even in early postnatal development ^13–15^. However, neural activity in sensory systems is principally driven by external stimuli and is thus sensitive to aberrations in upstream sensory processing as well as attentional control, each of which may also be affected in ASD/ID ^16,17^. These different influences highlight the need to develop new paradigms to accurately assess the functional properties of cognitive circuits in juvenile rat models of ASD/ID, both when such activity is driven by and when it is independent of external stimuli.

In contrast to primary sensory systems, neurons in cortical networks that give rise to the brain’s cognitive map often exhibit narrow tuning related to the animal’s current position in space ^18^. This spatial selectivity enables detailed mapping of their receptive fields in juvenile rats engaged in simple spatial exploration ^19–22^. Importantly, neural activity in these networks is constrained to low-dimensional neural manifolds that persist across online (e.g. awake) and offline (e.g. sleep) states ^23,24^. Recent work has shown that such manifolds provide access to neuronal tuning independently of the stimulus space ^23–27^, which makes these circuits ideal candidate model systems to study early postnatal network development *in vivo*.

The rodent head-direction (HD) circuit is a canonical example of a low-dimensional neural system in which population activity represents a one-dimensional circular variable ^28^. In the awake state, this latent variable corresponds to the animal’s current HD in the allocentric reference frame, while during sleep it no longer relates to the external world and instead represents an internally generated and drifting ‘manifold HD’ ^23^. This invariance of network topology across brain states indicates rigid functional connectivity, which can be probed by examining the neural manifold and the inferred latent variable.

Here, we show that in a rat model of Fragile X Syndrome (FXS), an inherited monogenic cause of ASD/ID ^29^, the cortical HD signal is transiently enhanced early in postnatal development but becomes unstable by adulthood. Using manifold analysis across online and offline brain states, we establish that the juvenile enhancement is largely due to differences in intrinsic dynamics while deterioration during adulthood stems from the difficulty with anchoring the HD signal to the external world. Overall, we introduce the rodent HD circuit as a valid and tractable model system that allows for unprecedented insights into network-level phenotypes in rodent models of neurodevelopmental disorders.

## Results

We recorded populations of neurons in the postsubiculum (PoSub), the cortical hub of the HD system ^30^, in juvenile post-weaning (P22-28) male rats missing the only copy of the *FMR1* gene (*Fmr1^−/y^* rats; 372 units in 5 rats) as well as in their wild-type (WT) littermates (406 units in 7 rats). PoSub cells were recorded using silicon probes (Figure 1a, Figure S1a-b) while rats foraged for scattered cereal in a cylindrical open field (Figure 1a). Each recording day consisted of two 20 min foraging sessions, with a 90 min period of rest between sessions, during which sleep activity was recorded. Juvenile *Fmr1^−/y^* rats exhibited similar exploratory behaviour (Figure S1b-f) and movement speed (Figure S1e) as their WT littermates. However, they spent significantly less time facing the polarizing visual landmark (Figure S1e).

**Figure 1.**
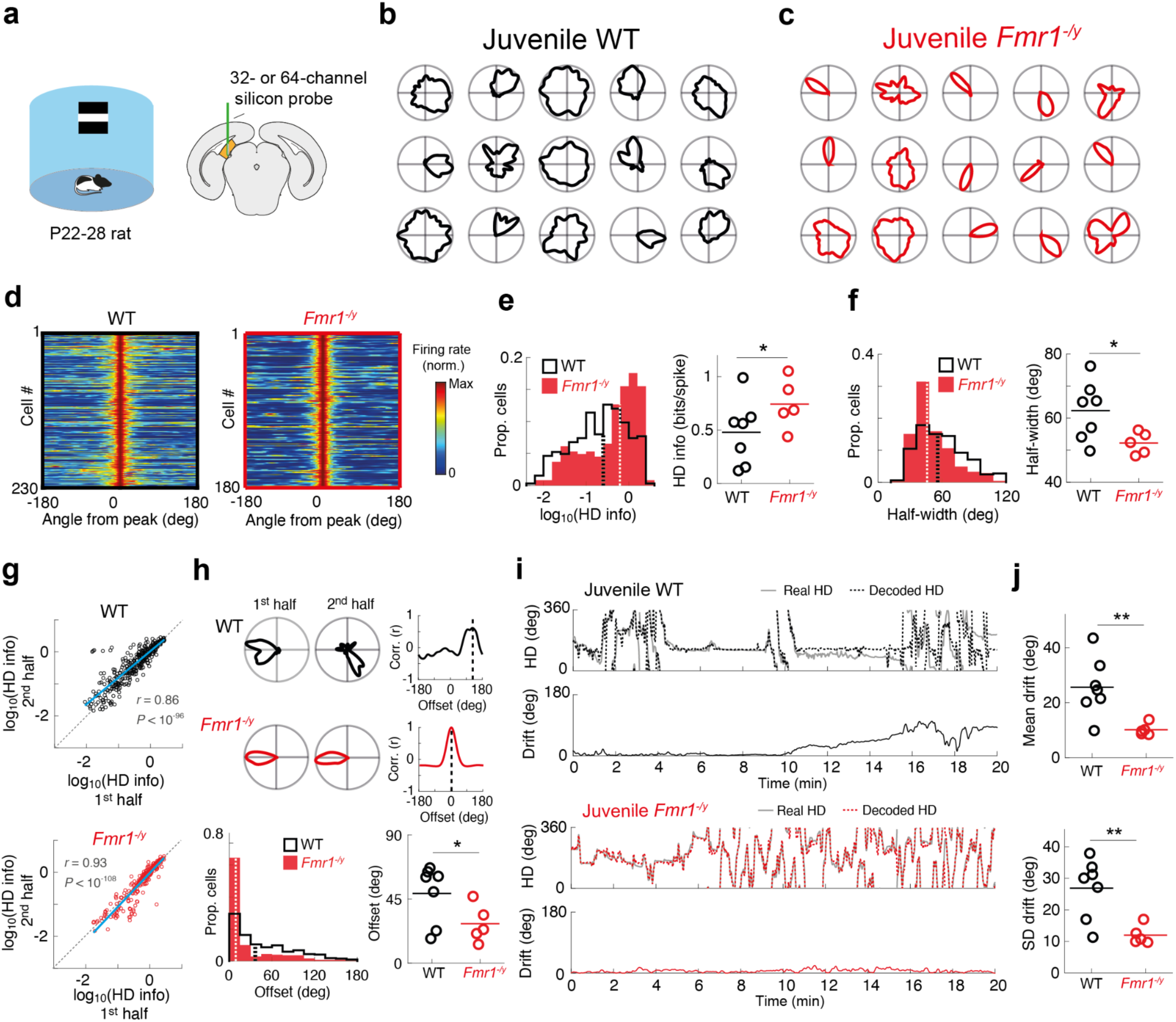
Enhanced HD tuning and HD signal stability in juvenile *Fmr1^−/y^* rats. **a**, Recording enclosure and silicon probe placement in PoSub. **b**,**c**, Example tuning curves of a subset of PoSub cells in a juvenile WT rat (**b**) and a juvenile *Fmr1^−/y^* rat (**c**). **d**, Normalized HD tuning curves of all PoSub cells with tuning curve width of less than 120 degrees. **e**, **f**, Overall distribution (left) and animal averages (right) of HD info (**e**) and tuning curve width (**f**) indicate enhanced HD tuning in juvenile *Fmr1^−/y^* rats (HD info, LME: genotype effect, *P* < 0.05; half-width, LME: genotype effect, *P* < 0.05). **g**, Relationship between HD tuning of PoSub cells across two halves of the exploration epoch (WT, *r* = 0.86, *P* < 10^−96^; *Fmr1^−/y^*, *r* = 0.93, *P* < 10^−106^). Blue lines, linear fits. **h**, Top, examples of HD tuning curves for two halves of a recording epoch and associated tuning cross-correlograms, showing considerable drift in a PoSub cell from a WT rat but a stable firing direction in a cell from an *Fmr1^−/y^* rat. Dashed lines, angular offset of maximum tuning curve correlation. Bottom, distribution of tuning curve offsets between two halves of the exploration epoch for all cells, as well as animal averages, indicating enhanced anchoring (i.e. smaller offsets) of the HD signal to the allocentric reference frame in *Fmr1^−/y^* rats (angular offset between tuning curves, LME: genotype effect, *P* < 0.05). **i**, Example real and decoded HD during a single recording session in WT and *Fmr1^−/y^* rats, as well as the drift of the decoded HD with respect to the animal’s real HD. **j**, Juvenile *Fmr1^−/y^* rats show less drift of the HD signal within the recording session (t-test on animal averages: mean drift, t(10) = 3.17, *P* < 0.01; Standard deviation (SD) of drift, t(10) = 3.37, *P* < 0.01). Dashed lines on histograms, population medians; horizontal lines on animal average panels, means of animal averages.

Excitatory cells in PoSub have been shown to be robustly tuned to HD in both adult and post-weaning juvenile WT rats ^21,31^, so we limited our analysis to this putative cell type, using trough-to-peak duration as the classification criterion (Figure S2a; range: 8-83 putative excitatory units per recording). While we observed directional tuning of putative excitatory cells in PoSub (henceforth referred to as PoSub cells) in both WT and *Fmr1^−/y^* juvenile rats, to our surprise well-tuned HD cells constituted a larger proportion of the recorded cell population in the latter (Figure 1b-d). Quantification of HD tuning using either via HD information (‘HD info’) or the width of the tuning curves (Figure 1e-f, Figure S2e-f), confirmed that overall, PoSub cells in juvenile *Fmr1^−/y^* rats were indeed more sharply tuned to HD than PoSub cells in their WT littermates. This difference in tuning could not be explained by differences in cluster quality, firing rates, electrode positions or age at the time of recording (Figure S2b-d, Figure S2g-h).

The narrower HD tuning curves of juvenile *Fmr1^−/y^* PoSub cells could be explained either by improved anchoring of the HD signal to the external world or by intrinsically enhanced HD tuning compared to WT. To assess the former explanation, we first compared HD tuning between the first and second half of each 20 min foraging session. While the robustness of HD tuning (head direction information) did not differ between the two halves of the recording session in either genotype (Figure 1g), the tuning curves in juvenile *Fmr1^−/y^* rats drifted less between the two halves of the session, with respect to the allocentric reference frame, than those of WT rats (Figure 1h). This improved anchoring of the HD signal to the external world in juvenile *Fmr1^−/y^* rats was further confirmed on the population level (Figure 1i-j). A similar pattern of results was observed when comparing HD stability between sessions, with greater stability in *Fmr1^−/y^* than WT rats (Figure S2i; Figure S21j). Thus, the apparent enhancement of HD tuning in juvenile *Fmr1^−/y^* rats could be explained, at least in part, by the lower degree of drift of the HD signal with respect to the allocentric reference frame.

If the differences in anchoring to the external world underlie the differences in HD tuning, we would expect the overall HD tuning in juvenile *Fmr1^−/y^*rats to be similar to that of WT rats if computed independently of the allocentric reference frame. In other words, the fidelity of the internally generated HD signal (the ‘manifold HD’) should be the same across the two groups, with the differences lying in how this internal HD signal relates to the animal’s real HD with respect to the environment. Highly structured intrinsic dynamics within the HD circuit enable estimation of relative HD tuning curves from population activity alone, independent of any external variables and reference frames ^23^. We thus applied the Isomap dimensionality reduction technique to the PoSub cell population vectors obtained during the exploration sessions (Figure S1a). In both WT and *Fmr1^−/y^* rats, the radial coordinate of PoSub population activity projected onto two dimensions (manifold HD) corresponded to the animal’s current HD and could be used to compute manifold HD tuning curves (Figure 2a-e). However, while HD tuning curves reconstructed from the Isomap projection corresponded well to the HD tuning curves calculated in the allocentric reference frame (Figure 2c-d, Figure 2f), these intrinsic tuning curves were better tuned to HD in juvenile *Fmr1^−/y^* rats than in their WT littermates (Figure 2g, Figure S2b-c). Thus, PoSub cells in juvenile *Fmr1^−/y^* rats exhibit intrinsically enhanced tuning to HD independently of the differences in anchoring to the allocentric reference frame.

**Figure 2.**
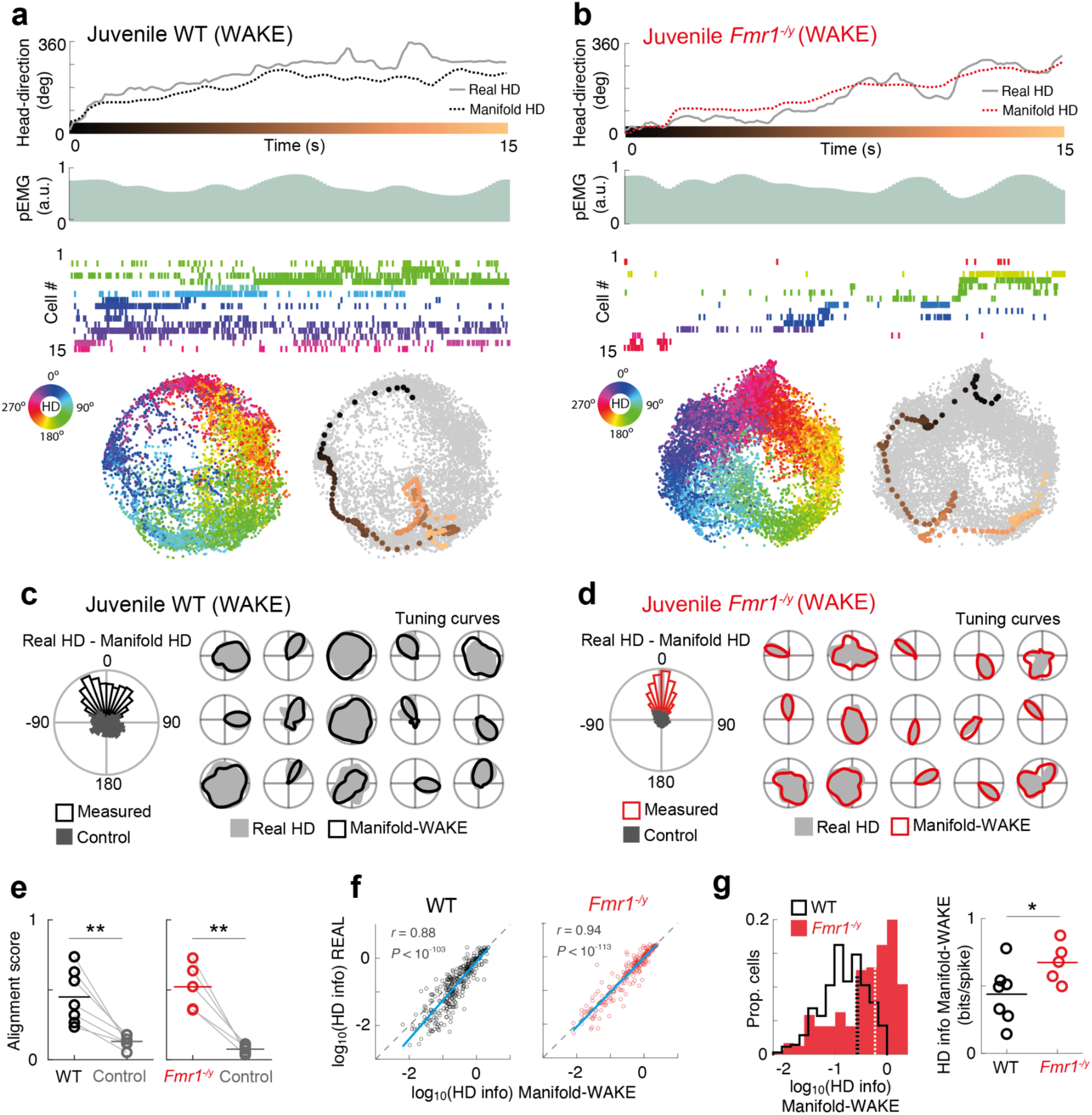
PoSub cells in juvenile *Fmr1^−/y^* rats show enhanced tuning to the internal HD signal during wakefulness. **a**, **b**, Examples of manifold HD (top) decoded from Isomap projections (bottom) of PoSub cell spiking (middle, 15 cells with highest HD info) during a single awake exploration session (WAKE) in a juvenile WT rat (**a**) and an *Fmr1^−/y^* rat (**b**), with each population vector colored according to the animal’s current HD. pEMG, pseudo-electromyogram. **c, d,** Alignment of the manifold HD (angle from the centre of the manifold) to the real HD measured externally (left) and example HD tuning curves computed in the allocentric reference frame (Real HD) and reconstructed using the angle from the centre of the manifold (Manifold-WAKE). **e,** In all animals alignment of the manifold HD with real HD was stronger than the control computed by reversing the manifold HD in time (WT: t(12) = 4.60, P < 0.01; *Fmr1^−/y^*: t(8) = 5.87, P < 0.01). **f**, Magnitude of HD tuning in PoSub cells is preserved in the tuning curves reconstructed only from the manifold during WAKE (WT, *r* = 0.88, *P* < 10^−103^; *Fmr1^−/y^*, *r* = 0.94, *P* < 10^−113^). Blue lines indicate linear fits. **g**, Overall distribution (left) and animal averages (right) of HD info indicate enhanced tuning to manifold HD in juvenile *Fmr1^−/y^* rats (LME: genotype effect, *P* < 0.05). Dashed lines on the histogram indicate population medians; horizontal lines on animal average panels are means of animal averages.

While the manifold HD obtained from population activity during wake periods should be independent of drift, it is still conceivable that it is to some degree influenced by ongoing behaviour. In contrast, population activity during sleep is internally generated and dictated by the internal dynamics. We thus took advantage of the fact that HD cell population dynamics during Rapid Eye Movement (REM) sleep are virtually indistinguishable from wakefulness, with the HD signal drifting at real-time speed despite the animal being stationary ^23,25,28^. Manifold HD can be reliably decoded from HD cell population activity during REM and HD tuning curves can be reconstructed purely from internally generated activity in absence of consciousness (Figure 3a-b). We found that, overall, juvenile *Fmr1^−/y^* rats spent a smaller proportion of sleep in REM (Figure 3c, Figure S3d-e), but the REM spiking activity was similar to wakefulness in both groups (Figure S3f). HD tuning curves generated from the REM manifold corresponded well to the HD tuning curves calculated in the allocentric reference frame during exploration but were free from drift (Figure 2d, Figure S3g-i). Importantly, PoSub cells in juvenile *Fmr1^−/y^* rats still showed enhanced HD tuning even when the tuning was computed from internally generated activity during REM sleep (Figure 2e). Overall, these findings indicate that on top of enhanced anchoring to the allocentric reference frame, HD tuning in juvenile *Fmr1^−/y^* rats is intrinsically more precise.

**Figure 3.**
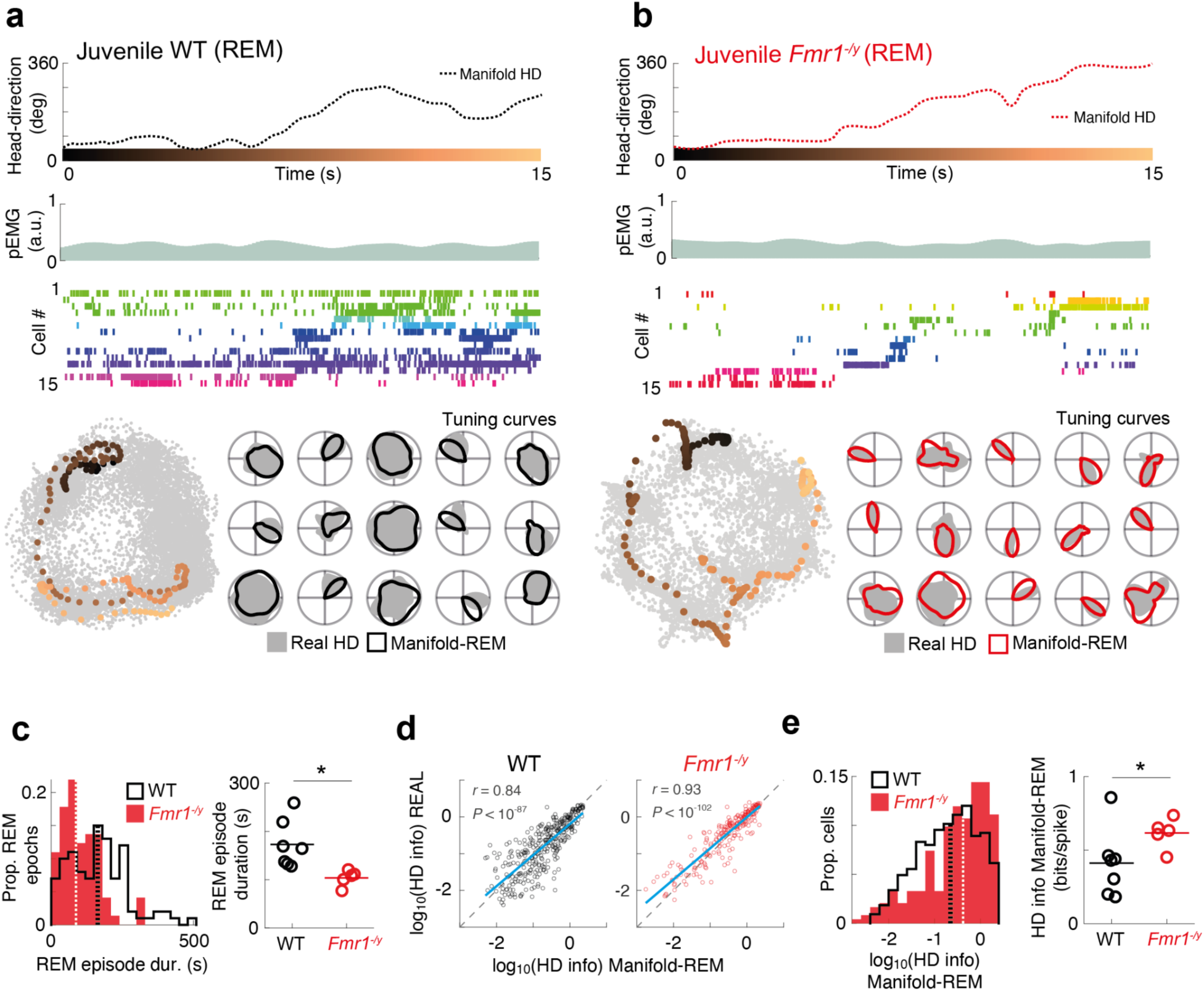
Enhanced PoSub cell tuning to the internal head direction signal in juvenile *Fmr1^−/y^* rats persists during REM sleep. **a**, **b**, Examples of manifold HD (top) decoded from Isomap projections (bottom left) of PoSub spiking activity (middle) during REM sleep from the same recording session as in Figure 2a-d. HD tuning curves (bottom right) can be reliably reconstructed from the REM activity (Manifold-REM), in the absence of consciousness (i.e. offline). **c**, Juvenile *Fmr1^−/y^* rats have shorter REM episodes (LME: genotype effect, *P* < 0.05). **d**, Magnitude of HD tuning in PoSub cells is preserved in the Isomap REM tuning curves in both genotypes (WT, *r* = 0.84, *P* < 10^−87^; *Fmr1^−/y^*, *r* = 0.93, *P* < 10^−102^). Blue lines, linear fits. **E**, Overall distribution (left) and animal averages (right) of HD info indicate enhanced HD tuning to internally generated activity during REM in juvenile *Fmr1^−/y^*rats (LME: genotype effect, *P* < 0.05). Dashed lines on histograms, population medians; horizontal lines on animal average panels, means of animal averages.

We then sought to establish whether the unanticipated enhancement in the fidelity of cortical HD signal in juvenile *Fmr1^−/y^* rats is transient or persists to adulthood. To that end, we recorded PoSub activity in adult *Fmr1^−/y^* rats (n = 372 units from 6 rats) and their WT littermates (n = 406 units from 6 rats) in the same experimental protocol (range: 30-114 putative excitatory units per recording; Figure 4a, Figure S4, Figure S5a-d). In contrast to what we observed in juvenile rats, HD tuning in adult *Fmr1^−/y^* rats was no different than that of WT littermates (Figure 4b-f; Figure S5e-g). However, while juvenile *Fmr1^−/y^* rats exhibited superior anchoring of the HD signal to the external world, adult *Fmr1^−/y^* rats showed an inverse phenotype, with more drift of the HD signal both within individual sessions (Figure 4g-j) and between the two recording epochs conducted 90 min apart (Figure S4h-i). As expected, the internal HD generated from the manifolds matched real HD well (Figure 5a-b, Figure 5e) and there were no differences in the tuning of HD tuning curves obtained from the manifolds during periods of wakefulness (Figure 5a-d). When we extended our investigation to internally generated activity during sleep (Figure 6a-b), we found that the REM sleep deficit observed in juvenile rats was ameliorated upon reaching adulthood (Figure 6c, Figure S6d-f). While the HD tuning curves generated from REM activity matched those generated from real HD (Figure 6d, Figure S6g), we did not observe any differences in such internally generated HD tuning between adult WT and *Fmr1^−/y^* rats (Figure 6e, Figure S6h-i). Thus, the enhancement in the quality of the intrinsic HD receptive fields in *Fmr1^−/y^* rats is developmentally transient and disappears as the rats reach adulthood. Furthermore, the superior anchoring of the HD signal to the external world observed in juvenile rats seems to degrade in adulthood, resulting in significant drift of the HD signal with respect to the allocentric reference frame despite intact intrinsic HD tuning.

**Figure 4.**
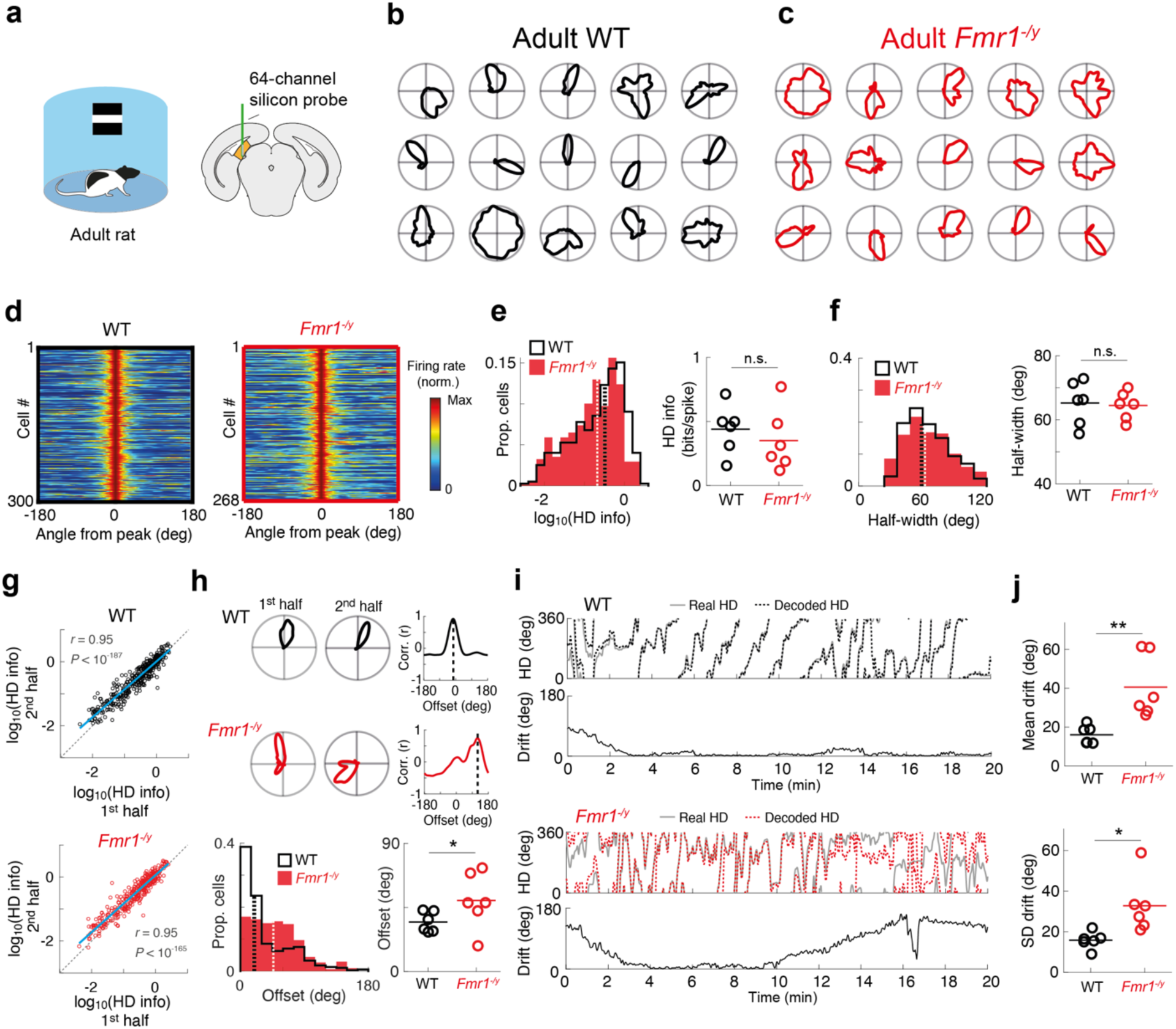
WT-like HD tuning but degraded HD signal stability in adult *Fmr1^−/y^* rats. **a**, Recording enclosure and silicon probe placement in PoSub. **b**,**c**, Example tuning curves of PoSub cells in adult WT (**b**) and *Fmr1^−/y^* rats (**c**). **d**, Normalized HD tuning curves of PoSub cells with tuning curve width of less than 120 deg. **e**, **f**, Overall distribution (left) and animal averages (right) of HD info (**e**) and tuning curve width (**f**) indicate normal levels of overall HD tuning in adults *Fmr1^−/y^*rats (HD info, LME: no genotype effect, *P* > 0.05; half-width, LME: no genotype effect, *P* > 0.05). **g**, Degree of HD tuning in PoSub cell tuning curves is preserved between two halves of an exploration epoch, indicating stable tuning fidelity (WT, *r* = 0.95, *P* < 10^−187^; *Fmr1^−/y^*, *r* = 0.95, *P* < 10^−165^). Blue lines indicate linear fits. **h**, Top, examples of HD tuning curves for two halves of a recording epoch and associated tuning cross-correlograms, showing considerable drift in a PoSub cell from an *Fmr1^−/y^* rat but stable firing direction for a PoSub cell in a WT rat. Dashed lines indicate the angular offset of maximum tuning curve correlation. Bottom, overall distribution tuning curve offsets between two halves of the exploration epoch for all cells as well as animal averages (angular offset between tuning curves, LME: genotype effect, *P* < 0.05). **i**, Example real and decoded HD during a single recording session in WT and *Fmr1^−/y^* rats, as well as the drift of the decoded HD with respect to the animal’s real HD. **j**, Adult *Fmr1^−/y^* rats show increased drift of the HD signal within the recording session (t-test on animal averages: mean drift, t(10) = 3.55, *P* < 0.05; SD drift, t(10) = 2.90, *P* < 0.01). Dashed lines on histograms are population medians; horizontal lines on animal average panels indicate means of animal averages.

**Figure 5.**
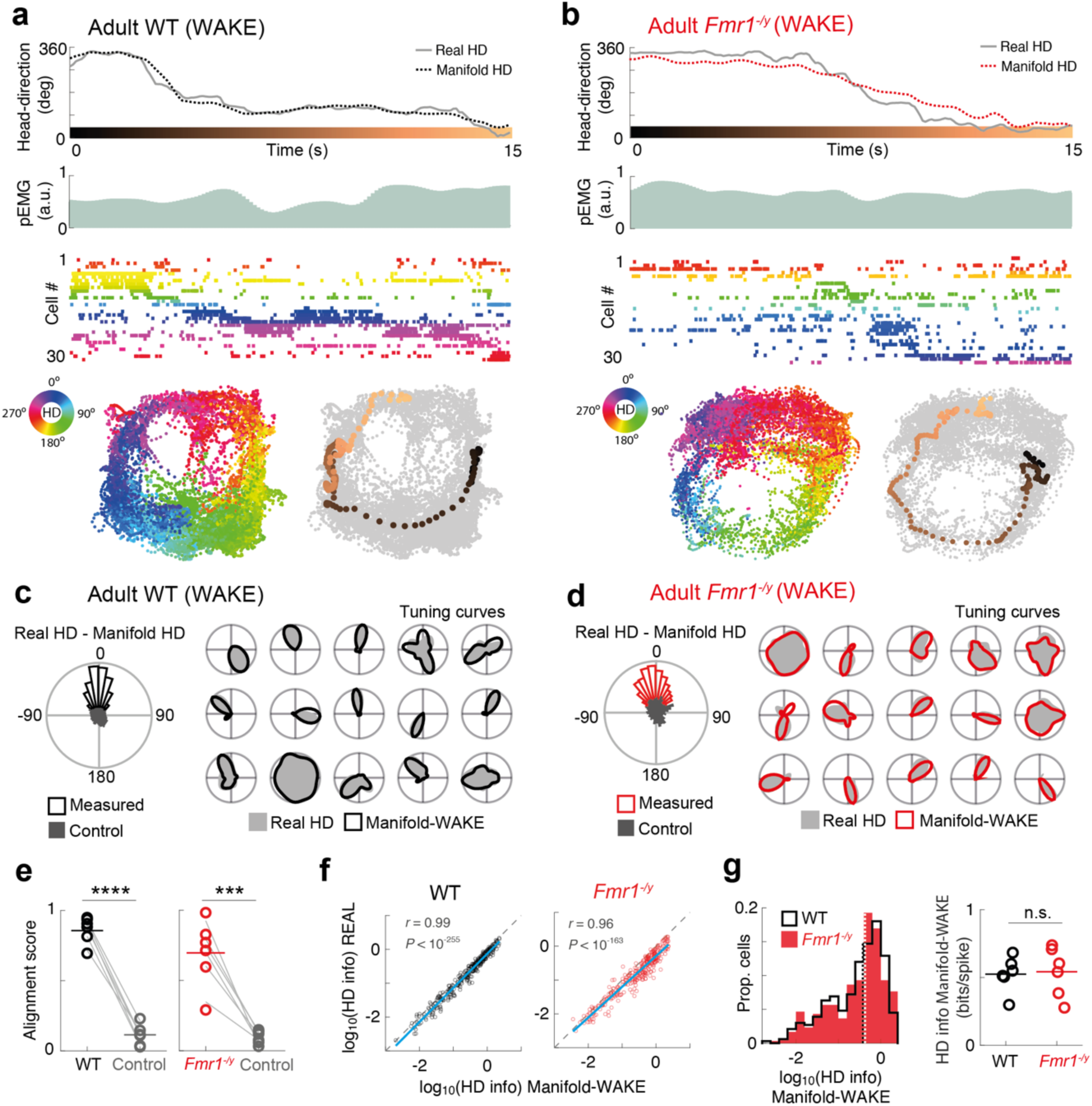
PoSub cells in adult *Fmr1^−/y^* rats show WT-like tuning to the internally generated HD signal during wakefulness. **a**, **b**, Examples manifold HD (top) decoded from Isomap projections (bottom) of PoSub cell spiking (middle, 30 cells with highest HD information) during a single awake exploration session (WAKE) in an adult WT rat (**a**) and an *Fmr1^−/y^* rat (**b**), with each population vector colored according to the animal’s current HD. pEMG, pseudo-electromyogram. **c, d,** Alignment of the manifold HD to the real HD measured externally (left) and example HD tuning curves computed in the allocentric reference frame (Real HD) and reconstructed using the angle from the centre of the manifold (Manifold-WAKE). **e**, In all animals, alignment of the manifold HD with real HD was stronger than the control computed by reversing the manifold HD in time (WT: t(10) = 19.6, P < 10^−5^; *Fmr1^−/y^*: t(10) = 7.13, P < 0.001). **f**, HD tuning in PoSub cells is preserved in manifold HD tuning curves during WAKE (WT, *r* = 0.99, *P* < 10^−255^; *Fmr1^−/y^*, *r* = 0.96, *P* < 10^−163^). Blue lines indicate linear fits. **g**, Overall distribution (left) and animal averages (right) of HD info indicate WT-like HD tuning in adult *Fmr1^−/y^* rats when measured independently of any reference frames (LME: no genotype effect, *P* > 0.05). Dashed lines on histograms are population medians; horizontal lines on animal average panels indicate means of animal averages.

**Figure 6.**
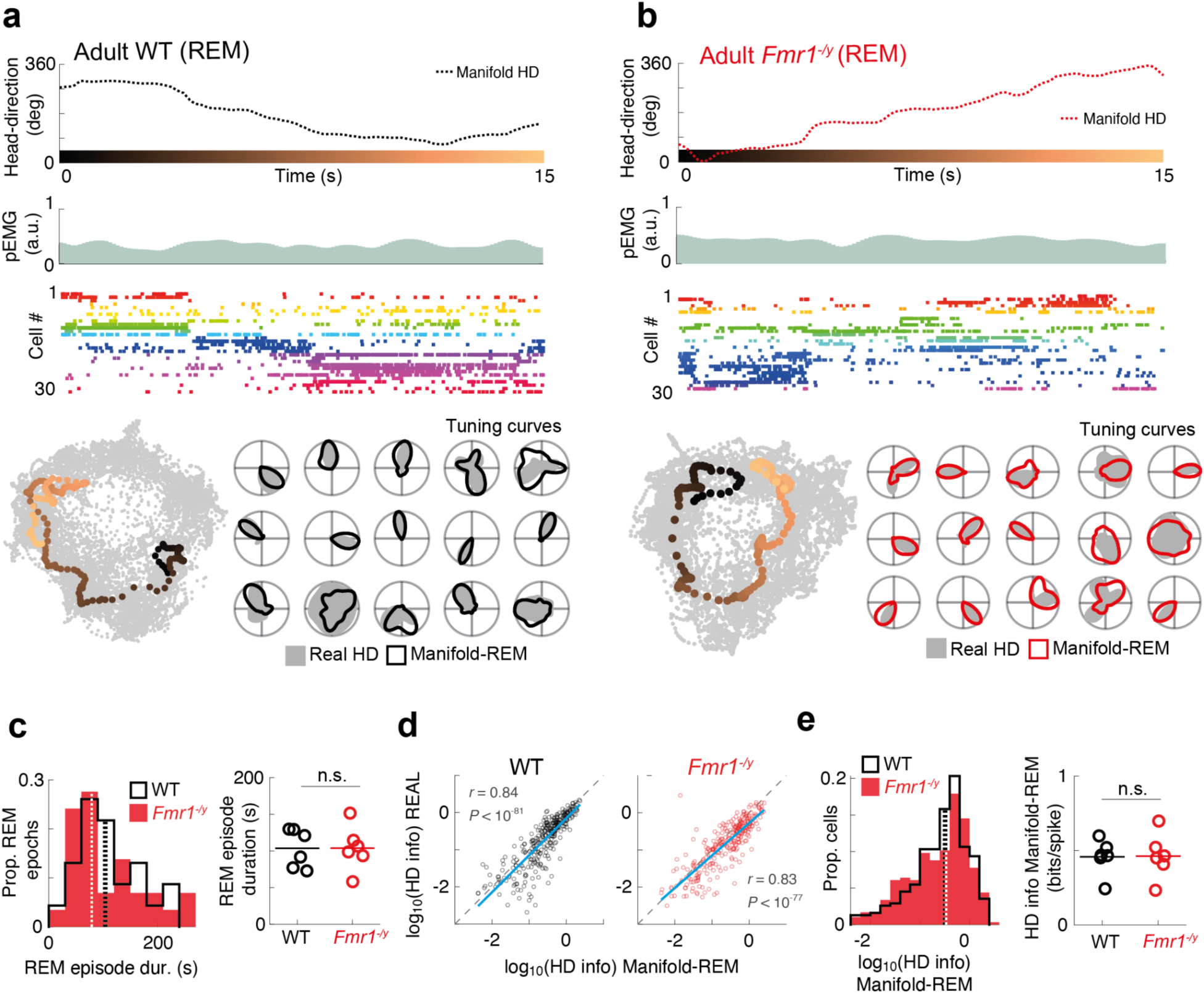
PoSub cells in adult *Fmr1^−/y^* rats show WT-like tuning to the internally generated HD signal during REM sleep. **a**, **b**, Examples of manifold HD (top) decoded from Isomap projections (bottom left) of PoSub spiking activity (middle) during REM sleep from the same recording session as in Figure 4a-d. HD tuning curves (bottom right) can be reliably reconstructed from the REM activity (Manifold-REM), in the absence of consciousness (i.e. offline). **c**, Adult *Fmr1^−/y^* rats have REM episodes of WT-like duration (LME: no genotype effect, *P* > 0.05). **d**, Magnitude of HD tuning in PoSub cells is preserved in the Isomap REM tuning curves in both genotypes (WT, *r* = 0.84, *P* < 10^−81^; *Fmr1^−/y^*, *r* = 0.83, *P* < 10^−77^). Blue lines, linear fits. **e**, overall distribution (left) and animal averages (right) of HD info indicate WT-like HD tuning to internally generated activity during REM in adult *Fmr1^−/y^*rats (LME: no genotype effect, *P* > 0.05). Dashed lines on histograms, population medians; horizontal lines on animal average panels, means of animal averages.

## Discussion

In the present study, we assessed how neural representations of the outside world are altered in a rat model of Fragile X Syndrome. We leveraged the experimental tractability of the cortical head direction cell system in *Fmr1^−/y^* rats to reveal an unexpected enhancement of HD tuning in PoSub that is apparent in 3-4 week old juveniles but disappears by adulthood. This phenotype was accompanied by an enhanced anchoring of the HD signal to the external world, but reconstruction of the intrinsic HD tuning curves from PoSub cell activity manifolds during both wakefulness and REM sleep allowed us to show that the enhanced fidelity of the HD signal in juvenile *Fmr1^−/y^* rats was independent of anchoring. In adult *Fmr1^−/y^* rats, in contrast, the anchoring of the HD signal to external landmarks was weaker than that of WT rats. Thus, our results reveal a developmental regression in this neural representation in *Fmr1^−/y^* rats.

Previous spatial phenotypes reported in *Fmr1^−/y^* rats have been subtle, and highly dependent on age, brain area, and experimental conditions. Impairments in spatial memory and cognition have been reported in *Fmr1^−/y^* rats, including slower and poorer performance in a Morris water maze task ^32^ and a deficit in object-place-context recognition memory ^3,4^, but both of these phenotypes were sensitive to conditions such as experimental age and the exact training paradigm. Within the hippocampus, *Fmr1^−/y^* neurons showed reduced experience-dependent changes in activity ^5^. Consistent with individuals with ASD, *Fmr1^−/y^* rats show differences in sensory processing compared to wild type rats. This includes degraded responses to auditory stimuli ^33^, faster reaction times to sound and impaired temporal but enhanced spectral modulation of loudness, compared to control ^34^, inappropriate hyperactivity in the juvenile visual cortex ^35^, and altered neurological and behavioural responses to almond odour ^36^. Thus, the phenotypes reported here are consistent with a pattern of neuronal alterations in spatial cognition and sensory processing.

The enhanced head direction tuning in juvenile *Fmr1^−/y^* rats may be local to the PoSub, or be a product of alterations earlier in the HD circuity. The head direction signal is thought to originate in the network comprising the dorsal tegmental nucleus (DTN) in the brainstem and the lateral mammillary nucleus (LMN), which in turn projects via the anterior nucleus of the thalamus to PoSub ^37,38^. Modeling ^39–42^ and lesion ^43,44^ studies indicate that directional firing fields of HD cells are a consequence of a highly structured connectivity pattern in the DTN-LMN network, whereby cells coding for similar directions inhibit cells that fire for opposite directions, likely via inhibitory interneurons. Such a ring attractor network is constrained to the same state space independently of brain state, which allows for a coherent HD signal to be decoded as a latent variable during sleep ^25,28,45,46^. The exact connectivity within this network determines the precision of the latent variable and thus the tuning of individual HD cells. Thus, one possibility is that the enhancement of cortical HD tuning observed in juvenile *Fmr1^−/y^* rats is a consequence of altered connectivity in the upstream DTN-LMN circuit.

In contrast, the second main finding of the current work - the enhanced anchoring of the HD signal to the external world in juvenile *Fmr1^−/y^* rats - may arise from top-down cortical connections to the PoSub ^47^. Vision is the primary modality for this process, and the information about stable visual landmarks may reach PoSub either directly from the visual cortex, area 29e ^48^, or via prominent inputs from the retrosplenial cortex, where the activity of a fraction of neurons is selective for visual landmarks ^49,50^. Our data are consistent with altered processing in the pathway that conveys the visual landmark information to the HD circuit, but they do not preclude the possibility that juvenile *Fmr1^−/y^* rats enhance their anchoring by more efficient use of the olfactory ^51,52^, or somatosensory ^22^ systems.

The independence of HD directional tuning and anchoring to landmarks is further supported by our finding that in adult *Fmr1^−/y^* rats the intrinsic HD signal is intact across both wakefulness and sleep, but its anchoring to the external world is disrupted. An intact HD signal that is not correctly anchored to the allocentric reference frame is present in very young juvenile rats (P10-14) prior to eye opening ^22^, and in adult rodents it detaches from the external world whenever the visual inputs are compromised ^51^. In P12-14 rats, this anchoring deficit can be mitigated by providing polarising somatosensory information. It is thus possible that adult *Fmr1^−/y^* rats have a sensory deficit that interferes with the correct mapping of the manifold HD to the allocentric reference frame. Such deficit could include alterations in processing or integration of external visual or olfactory information and/or processing or integration of self-motion cues, such as angular velocity, optic flow and motor efference copy. All of these input streams may be used to update the HD signal as the animal moves around within the environment ^37,51,53^.

It is possible that phenotypes reported in *Fmr1^−/y^* rats may not be directly caused by the lack of FMRP; rather, neuronal alterations and the associated functional effects may be secondary adaptations to the loss of FMRP. For example, the hyperexcitability seen in the *Fmr1^−/y^* CA1 cells is associated with reduced functional inputs from the entorhinal cortex, meaning that it may result from adaptive or homeostatic regulation induced by reduced functional synaptic connectivity ^54^. Using ex vivo electrophysiology and computational modelling ^55^ showed that an increased excitation/inhibition ratio in the *Fmr1^−/y^* mouse somatosensory cortex does not lead to changed firing rates downstream, because altered excitatory and inhibitory synaptic conductances counteract each other. The result in vivo is a largely normal sensory tuning with a mildly blurred whisker map, suggesting that some of the changes in excitability seen in the animal models are appropriately tuned to counterbalance the direct effects of the mutation, to normalise the overall output as much as possible. Other studies have also confirmed that some later-emerging *Fmr1^−/y^* phenotypes arise as secondary adaptations to the direct effects of FMRP’s absence on the level of cellular excitability ^11,56^. In support of the possibility of hyperexcitability arising secondary to previous phenotypes, it has been shown that hyperexcitability sometimes arises later in development. In *Fmr1^−/y^* rats before eye-opening, responses in the visual cortex had reduced spike rates (hypoexcitability); after the fourth postnatal week, mild hyperexcitability emerged as increase in visually-evoked firing of putative excitatory neurons and reduced firing of putative inhibitory neurons ^57^. This may also explain why cellular and neuronal hyperexcitability is not found in every brain region and cell type, as each one may develop compensatory changes differently. This may be tied to the phenomenon of developmental regression, sometimes observed in children with autism, where already acquired skills are lost, usually around the 2nd and 3rd year of life ^58^. The deficient anchoring of the HD signal to the external world, reported here, emerged during later postnatal development, and therefore may arise as imperfect homeostatic adaptations to a lack of FMRP.

Finally, one of our incidental findings was that juvenile *Fmr1^−/y^* rats have more fragmented REM sleep. Juvenile rats have longer REM bouts compared to adults ^59^ and REM sleep is associated with learning ^60^ and may direct the course of brain maturation through the control of neural activity ^61^. In the cortex, REM sleep is responsible for crucial developmental processes such as pruning and strengthening dendritic spines appropriately ^62^, and experimental suppression of REM sleep alters behavioural, anatomical, and biochemical development ^61,63,64^. The finding of shorter REM bouts in juvenile *Fmr1^−/y^* rats is consistent with studies in human children with FXS, reporting disrupted sleep microstructure, less time in REM sleep, and frequent night awakenings ^65^, while more problematic health or behavioural characteristics were associated with higher likelihood of sleep problems ^66^. Thus, a more fragmented sleep with shorter REM bouts, especially during development, can be both an effect of lack of FMRP and a cause for some of the intellectual disability-related phenotypes observed in *Fmr1*-knockout animals and individuals with FXS.

In summary, in the current study we have shown that the head direction system is altered in *Fmr1^−/y^* rats. The nature of this alteration suggests an apparent early gain-of-function followed by a decline. This pattern is intriguing as it echoes developmental regression, in which autistic children may acquire certain cognitive skills early in life but later lose them. Additionally, while our focus here has been on the head direction cell system because of its tractability, the current results may imply alterations in other spatial representations in the brain, for example those provided by place cells and grid cells, which likewise provide internal representations of space that are anchored to external landmarks.

## Methods

### Animals

All experimental procedures were carried out under a UK Home Office project licence, approved by the Animal Welfare and Ethical Review Board (AWERB) of the University of Edinburgh, and conformed with the UK Animals (Scientific Procedures) Act 1986. Long-Evans male *Fmr1^−/y^* and WT rat littermates were bred in-house by crossing WT sires with *Fmr1*-heterozygous dams. The original null mutants were generated using zinc finger nucleases targeting *Fmr1* exon 1 with a construct containing the sequence coding for eGFP. The *Fmr1^−/y^* males from the resulting line do not express either FMRP or eGFP, as described by ^4^. The rats used for the current experiments were bred in-house and were housed with their parents and siblings up until weaning (P21). After weaning, animals were housed in mixed genotype littermate cages. They were kept on a 12h/12h light/dark cycle (lights on at 7am) and fed *ad libitum* on standard laboratory chow, with supplementation of water-soaked breeding chow after the surgery. 24 hours after the end of the surgery, adult rats were placed on mild food restriction to encourage foraging, with their body weight not falling below 90% of their free feeding weight. Rats were randomly allocated to the experiment from the litter. Experimenters were blind to the genotype of the animals throughout experiments including surgery, recordings, and pre-processing, and were unblinded for data analysis.

### Surgery

Rats were anaesthetised using 1.5-3.5% isoflurane, while the animal’s respiratory rate and heart rate were monitored using a PhysioSuite (Kent Scientific) and their body temperature was maintained at 37° using a homeothermic blanket system (Harvard Apparatus). 0.08ml per kg bodyweight of rimadyl (50mg/ml carprofen) was administered subcutaneously shortly after induction of anaesthesia. To secure the headcap and implant on the head, the skull was covered with dental adhesive (Optibond Universal). In adults, this was supplemented with three to five self-tapping jeweller’s screws. A silver ground wire was placed through the craniotomy directly above the cerebellum in juveniles or attached to one of the skull screws placed above the cerebellum in adult rats. A craniotomy was drilled over the left postsubiculum, and a silicon probe (Cambridge Neurotech, models H5, H6b or H7) was lowered into the brain, slightly above the target area (juveniles, from Bregma: anterior-posterior (AP), −6.30 mm; medial-lateral (ML), 3.10 mm; dorsal-ventral (DV), −1.60 mm; adults, from Bregma: AP, −7.64 mm; ML, 3.30 mm; DV, −1.70 mm). The silicon probe was mounted on a drive (3DNeuro or made in-house), secured onto the skull’s surface using dental acrylic. After surgery, the animals were allowed to recover on a heated bench (30°C) for at least one hour and returned to the animal house for at least 24 hours before recording. Juvenile rats were single housed after surgery and adults were single-housed or housed with a sibling in a cage with a divider separating them. Juvenile rats were implanted between P20 and P23 and adult rats were at least 7 weeks old at the time of implantation.

### Recording

The probes were initially implanted in the white matter above PoSub and were subsequently advanced slowly into PoSub (speed: 0.28 mm/h in 0.035 mm increments) 24-48h after implantation. The tissue was allowed to settle for at least 3h before recordings commenced. The recording environment consisted of a blue plastic cylindrical arena of 73 cm diameter, with 54 cm tall walls. A prominent visual cue card with vertical black and white stripes was positioned near the top of the wall. The rat was taken within the curtains surrounding the recording enclosure and plugged in. It was then placed within the cylindrical arena from the side opposite to the visual cue, cereals were scattered on the floor and the rat was left to forage and explore for 20 min. After that, the rat was given a 90-minute sleep opportunity in an opaque container placed inside the recording arena, during which recordings continued. The arena was swept and cleaned with chlorhexidine (Vetasept) spray between the exploration and sleep sessions.

For recording, the implanted silicon probe was connected to a headstage (RHD, Intan technologies). The headstage was connected to an Intan RHD standard SPI cable, which was connected to a 3D-printed commutator in the ceiling (custom-made in house). Signal was acquired at 30 kHz with an OpenEphys data acquisition system. For tracking the rat’s position and head direction, a red and a green marker were attached on the right and left of the headstage. Video capturing and marker tracking were performed using Bonsai software ^67^ at 40 Hz. For the juvenile recordings, video images were acquired through a Logitech C930e camera. To synchronise the image with the electrophysiology data, an Arduino microcontroller was programmed to send a random sequence of pulses to both the OpenEphys system and a LED light within the frame of the camera. The times when the LED light was on were detected by Bonsai and synchronisation of the pulses in the light and the electrophysiology data was done offline. For the adult recordings, images were acquired through an acA1300-75gc Basler camera with a LMVZ4411 Kowa lens, positioned at the ceiling of the rig. This camera was connected to the OpenEphys board, sending a TTL pulse for each frame taken, allowing the synchronisation of the two streams of information.

### Perfusion and probe recovery

After the end of data gathering, the rats were anaesthetised with 5% isoflurane and injected intraperitoneally with a lethal dose of pentobarbitone (≥1ml for adults and ≥ 0.3ml for juvenile rats). After their reflexes were gone, they underwent transcardial perfusion first with 0.9% saline, and then with 4% formalin or paraformaldehyde solution. The rat was then decapitated and the brain extracted. The probe and drive complex was carefully recovered from the headcap. After recovery, the probe was dipped in and out of a 1:5 solution of Contrad 70 liquid detergent in distilled water heated to roughly 60°C for 5 minutes, twice. After that, the probe was left to soak in a 1-2% fresh solution of tergazyme detergent, and finally another round of dips in warm Contrad was performed. The probe was then rinsed with distilled water, and channel impedances were measured in 0.9% saline using the OpenEphys system. If the recording quality of the previous rat was good and the impedance values were satisfactory (most channels <0.5 MOhm), the probe was reimplanted in the next rat.

### Spike sorting and cell classification

Spikes were first split automatically into clusters using Kilosort 2.0 (Pachitariu et al., 2016). Manual curation was performed while visualising the automatically created clusters on Klusters software. At this stage, any cluster without a clear waveform and clear dip in the spike train autocorrelogram at the 0-1 ms time bin was classified as noise and cluster pairs with similar waveforms and a clear dip in their spike train cross-correlograms at the 0-1 ms time bin were merged. Viable units were first identified as units that had a waveform with negative deflection (criterion aiming to exclude spikes from fibers of passage). Next, a cluster contamination score was computed from the spike autocorrelograms, as a ratio of the amplitudes of the 0-1 ms bin and the maximum amplitude within 0-100 ms. Units with cluster contamination scores below 0.5 were excluded from the analysis. Next, putative excitatory cells were classified on the basis of trough-to-peak duration of their average waveforms. The threshold for classification was based on the trough of the bimodal distribution of trough-to-peak durations and was set at 0.5 ms for juvenile rats and 0.4 ms for adult rats. For each analysis, units with positive waveforms and units with less than 100 spikes fired in the analyzed epoch were excluded.

### HD tuning curves

The animal’s HD was calculated based on the line perpendicular to the line connecting the red and green markers attached to the headcap. Periods with more than 1 second without viable tracking data were excluded from the analysis. HD tuning curves were then computed as the mean firing rate of the cell in each angular bin. Tuning curves were calculated in bins of 1° (spikes/time) and smoothed with a Gaussian kernel of 6° SD. For Isomap analyses, tuning curves were computed in 6° bins (with 12° SD Gaussian kernel) to account for the relatively short duration of REM sleep episodes in adult rats. Speed was computed as the difference in the animal’s position at successive time points and was smoothed with a moving average window of 0.25 s. HD tuning curves were computed from epochs when the animal’s speed exceeded 1.5 cm/s.

Half-width of the HD tuning curves was computed by first finding the bin with the highest firing rate, and then averaging the two sides of the tuning curve. Half-width was then defined as twice the angular distance between the maximal firing rate and the half-maximal firing rate of the resulting half of the tuning curve. HD information contained in the tuning curves calculated for N angular bins as:

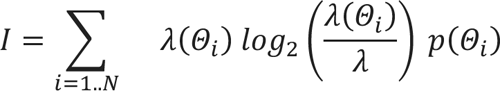

where *λ*(*Θ_i_*) is the firing rate of the cell for the *i*th angular bin and *λ* is the average firing rate of the neuron during exploration. We used uniform occupancy *p*(*Θ_i_*) to be able to compare tuning curve shapes irrespective of behavioural differences.

We obtained HD tuning curve crosscorrelograms by computing Pearson’s correlation coefficients between the reference tuning curve vector and the second tuning curve vector (from either the same or another cell), which was circularly shifted by 1 to maximum number of bins.

### Bayesian decoding and drift estimation

To estimate the degree of drift of the internal HD with respect to the allocentric reference frame, excitatory cell spike times were binned into population vectors (200 ms window, smoothed in 400 ms SD Gaussian windows). Based on cells’ tuning curves from the baseline period (first half of a recording epoch), the population vectors were converted into a Bayesian probabilistic map under the hypothesis that HD cells fire as a Poisson process ^68^. The instantaneous internal HDs were taken as the maxima of these probabilistic maps. The degree of drift of the HD signal was calculated as the decoder error: angular difference between the real HD and the decoded HD at each time bin. To obtain the slow drift signal, decoder error was denoised with a Hampel filter (5 s window, 1 SD) and then smoothened with a 0.8 s SD Gaussian window. The mean and SD of the decoding error was calculated from the second half of the recording session, restricted to times in which the animal’s velocity exceeded 1.5 cm/s. Recording sessions with less than 6 putative excitatory cells that fired more than 100 spikes were excluded from this analysis.

### Classification of sleep states

Sleep scoring was performed using the automated SleepScoreMaster algorithm and TheStateEditor software (Buzsaki laboratory, https://github.com/buzsakilab). The wide-band signal was downsampled to 1.25 kHz and used as the local field potential (LFP) signal. Pseudo-electromyograph (pEMG) was computed from correlated high-frequency noise across the channels of the linear probe.

### Visualization and analysis of the population manifold

For visualization of the PoSub manifold, cell spike times from the whole epoch were binned into population vectors in 200 ms bins, smoothened in 400 ms SD Gaussian windows and a square root of each value was taken. The resulting binned firing rates were then z-scored in the time domain and each population vector was then normalized by dividing it by its L2 norm. Population vectors were restricted to epochs in which animal’s velocity exceeded 1.5 cm/s (juvenile rats) or 3 cm/s (adult rats), and were then input into the Isometric Feature Mapping (Isomap) algorithm implemented in the Matlab Toolbox for Dimensionality Reduction Reduction (version 0.8.1b, https://lvdmaaten.github.io/drtoolbox/). The parameters were set to 25 nearest neighbours and 2 dimensions. Recording sessions with less than 6 putative excitatory cells that fired more than 100 spikes were excluded from this analysis.

Manifold HD at each time bin was then calculated as a four-quadrant arctangent of the two Isomap dimensions (range: −180° to 180°). The manifold HD generated this way has arbitrary directionality (clockwise/anticlockwise) and an arbitrary point of origin. To align the manifold HD obtained during wakefulness with real HD, manifold directionality was first established by computing the correlation between unwrapped real and manifold HD values and multiplying the manifold HD values by −1 if the correlation was negative. Manifold HD was then aligned to the real HD by computing the manifold offset as the circular mean of the difference between real and manifold HD, and circularly shifting all manifold HD values by this offset value. The alignment score was then computed as the Rayleigh vector length of all offset values.

Manifold tuning curves were then computed as the ratio between histograms of spike count and total time spent in each manifold HD in bins of 6 degrees and smoothed with a Gaussian kernel of 12° SD. Real HD tuning curves for comparison with manifold tuning curves were computed by downsampling the real HD into 200ms bins and applying the same procedure as above.

### Data analysis and statistics

All analyses were conducted using Matlab R2020b (Mathworks) with TStoolbox (https://github.com/PeyracheLab/TStoolbox) and Circular Statistics Toolbox version 1.2167 ^69^. Statistical comparisons of individual cell data were performed using a generalized linear mixed model (GLMM) approach ^70^ in order to take into account the hierarchy of dependency in our data sets (genotypes-rats-neurons) and account for random effects. Statistical modelling routines for the linear mixed effects (LME) models were written and run using RStudio 2023.12 (RStudio Team). Depending on the data distribution (normal, lognormal or gamma), linear mixed models (LMMs) or generalized linear mixed models (GLMMs) were fitted to single unit data metrics using the R package lme4 v1.1–17. Animal and cell identity were included in models as random effects, and genotypes were included as fixed effects. Terms are progressively eliminated when a simpler model (i.e. a model not containing that term) fits the data equally well (based on likelihood ratio test). Consequently, the p-values reported in the context of LMEs are given by likelihood ratio tests between a model containing the variable or interaction in question and a model without that variable or interaction (a reduced/null model). All statistically significant differences obtained through LMEs were also statistically significant when comparisons were made using a two-sample Mann-Whitney U test with cells as experimental units.

Statistical comparisons on animal averages (where stated) were performed with independent or paired Student’s t-tests. All statistical tests were two-tailed. No statistical methods were used to pre-determine sample sizes, but our sample sizes are similar to those reported in previous publications.

## Data Availability

Data will be made openly available at the time of publication.

## Code Availability

Code will be made openly available at the time of publication.

## Acknowledgements

NM was supported by a PhD studentship from the Simons Initiative for the Developing Brain. This work was supported by pilot funding from the Edinburgh-McGill pilot funding scheme (AP, ERW), and from the Simons Initiative for the Developing Brain (ERW & AA), the Simons Foundation Autism Research Initiative grant AN-AR-Rat Models-0090333 (AJD, PAD, and AP), the Canadian Research Chair in Systems Neuroscience (AP), CIHR Project Grants 155957 and 180330 (AP); NSERC Discovery Grant RGPIN-2018-04600 (AP). The authors thank Lynn Morrison and the other animal technicians from the University of Edinburgh George Square Bioveterinary Services Facility for their expertise and assistance in this work, and Owen Dando for help implementing the LME analysis.

---

**Figure S1.**
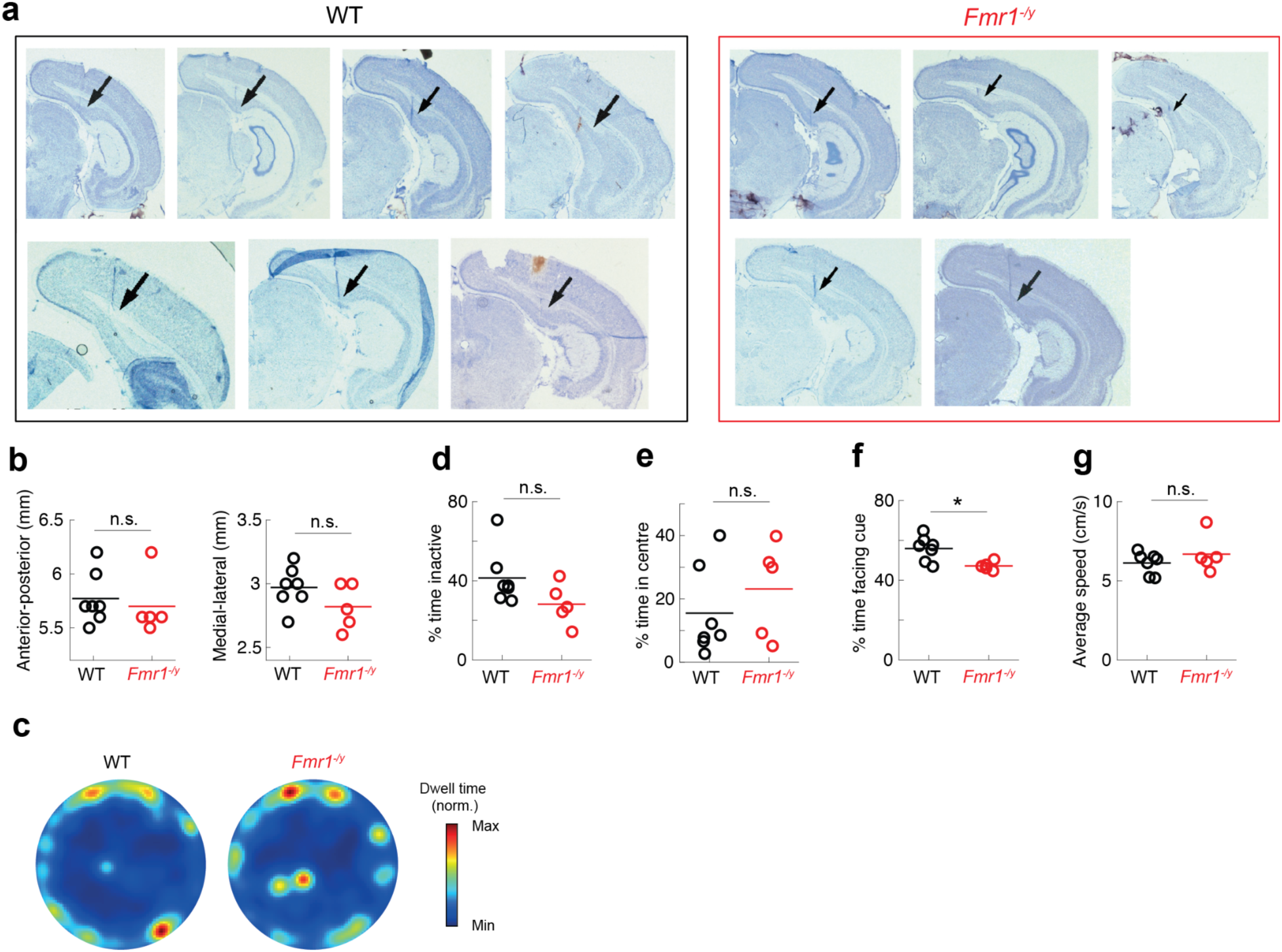
Probe placement and behaviour in juvenile WT and *Fmr1^−/y^*rats. **a**, Silicon probe tracts going through PoSub, as visualized by Nissl staining. Black arrows, electrode tracts. **b**, No significant differences in probe placement in the anterior-posterior (left) and medial-lateral (right) directions (t-test on animal averages, anterior-posterior: t(10) = 0.47, *P* > 0.05; medial-lateral: t(10) = 1.54, *P* > 0.05). **c**, Heat maps of average occupancy across the two genotypes. **d**, Both groups spent equal amounts of time inactive (t-test on animal averages: t(10) = 1.78, *P* > 0.05). **e**, Both groups spent the same amount of time in the centre of the arena (t-test on animal averages: t(10) = 0.90, *P* > 0.05). **f**, *Fmr1^−/y^* rats spent significantly less time than WT rats facing the visual landmark (t-test on animal averages: t(10) = 2.99, *P* < 0.05). **g**, WT and *Fmr1^−/y^* rats ambulated with the same average speed within the active period (t-test on animal averages: t(10) = 1.03, *P* > 0.05). Horizontal lines on animal average panels, means of animal averages.

**Figure S2.**
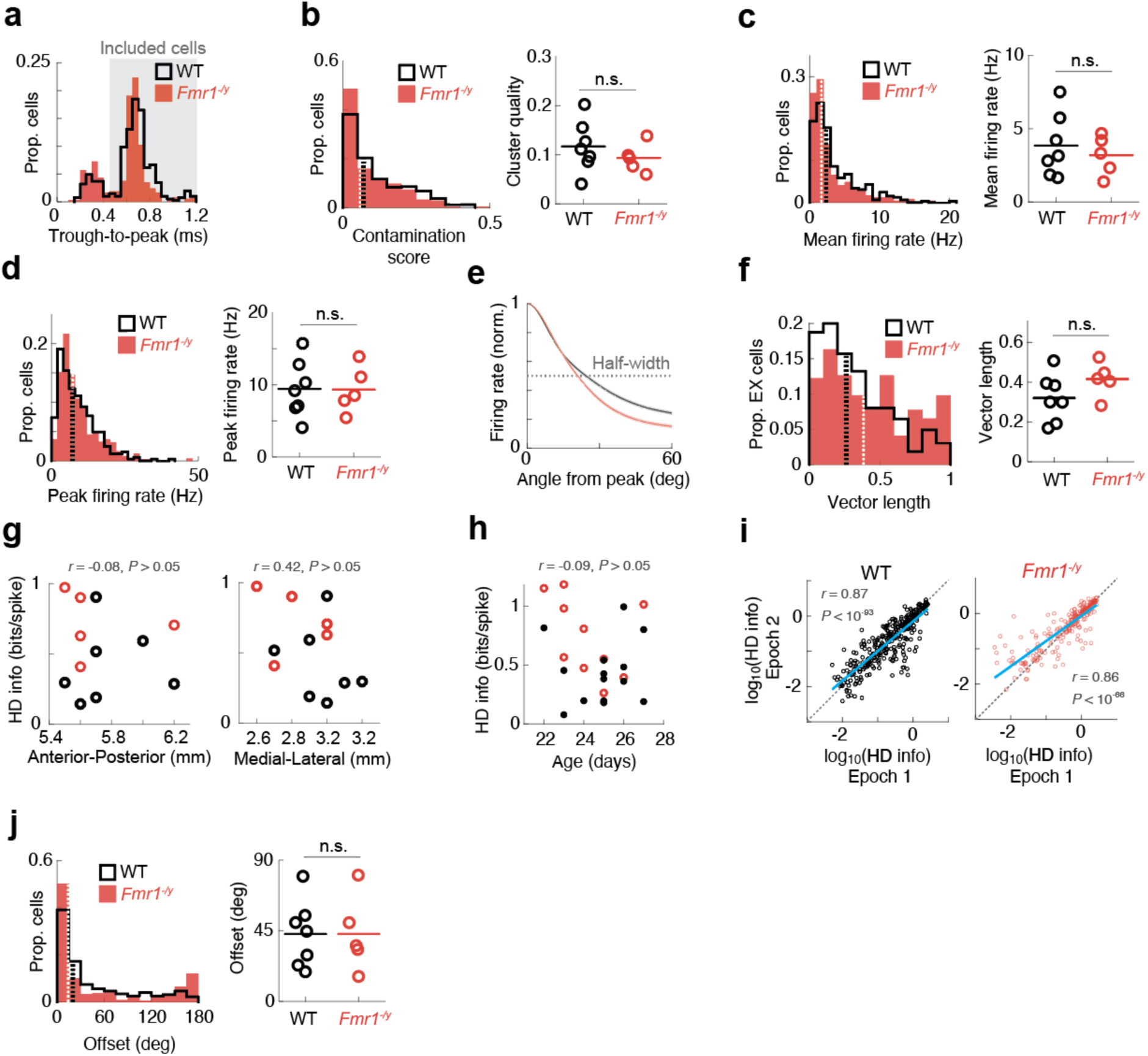
Single-cell and session metrics in juvenile WT and *Fmr1^−/y^* rats. **a**, Distribution of the trough-to-peak duration of all recorded units. Shaded area, cells classified as putative excitatory cells based on the trough in the bimodal distribution. **b**, Cluster contamination scores across the two groups (LME: no genotype effect, *P* > 0.05). **c**-**d**, Mean firing rates (LME: no genotype effect, *P* > 0.05) (**c**) and peak firing rates (**d**) of PoSub cells (LME: no genotype effect, *P* > 0.05). **e**, Mean normalized HD tuning curves of putative excitatory cells recorded in juvenile WT and *Fmr1^−/y^* rats. Only cells with tuning curve half-widths of less than 120 degrees were included. Shaded areas, +/- SEM. **f**, Resultant vector length across the genotypes (LME: no genotype effect, *P* > 0.05). **g**, No significant relationship between probe implantation coordinates and observed mean HD info for a given animal (anterior-posterior: r(10) = −0.08, P >0.05; medial-lateral: r(10) = −0.42, P > 0.05). **h**, No significant relationship between mean HD info for a given recording session and the age of the animal at the time of recording (r(22) = −0.09, P > 0.05). **i**, HD tuning is preserved across the two exploration epochs in both genotypes (WT, *r* = 0.87, *P* < 10^−93^; *Fmr1^−/y^*, *r* = 0.86, *P* < 10^−66^). **j**, Drift of HD cell receptive fields across the two recording epochs (LME: no genotype effect, *P* > 0.05). Dashed lines on histograms, population medians; horizontal lines on animal average panels, means of animal averages.

**Figure S3.**
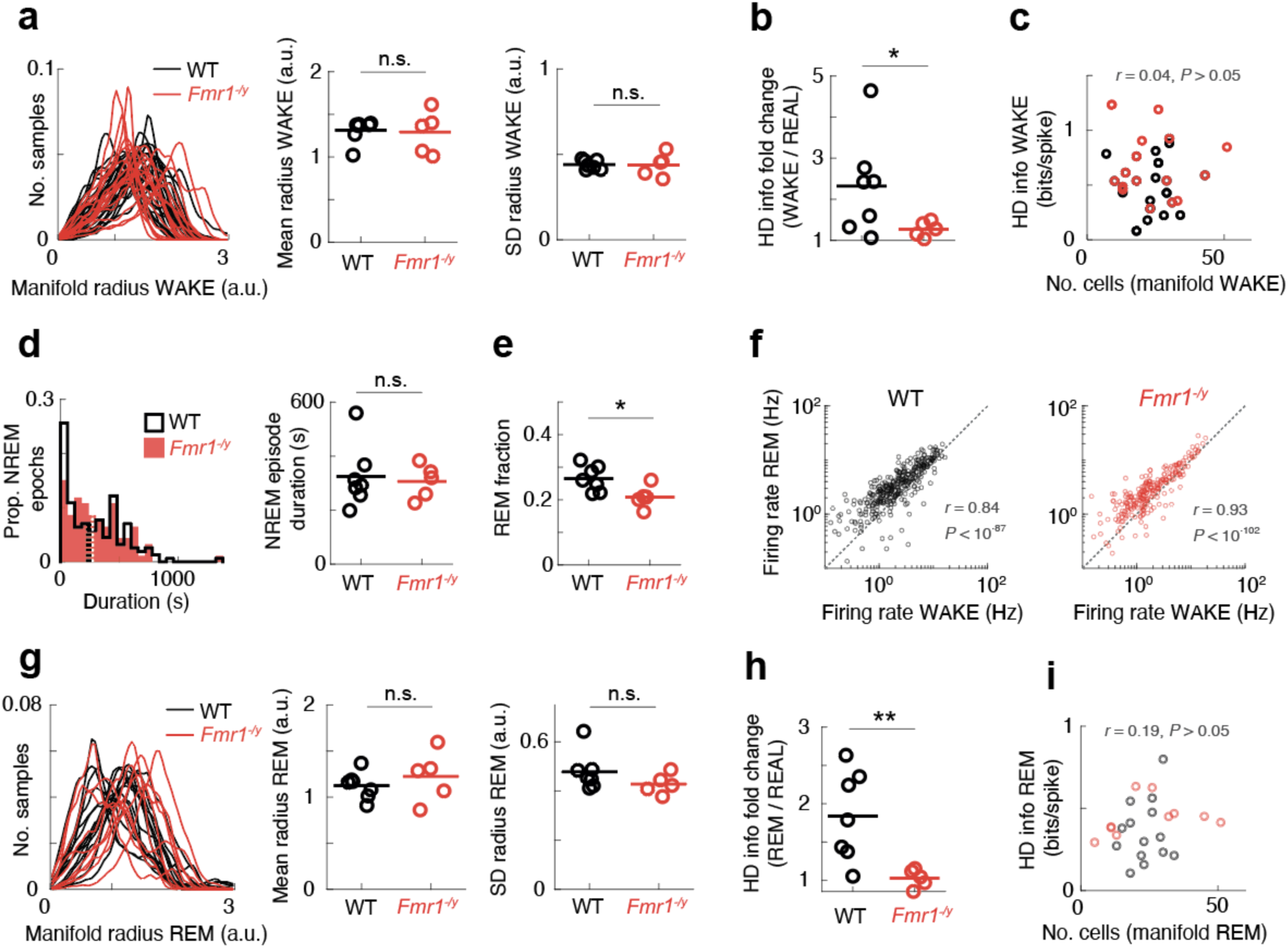
Manifold and sleep metrics in juvenile WT and *Fmr1^−/y^* rats. **a,** There were no significant differences between groups in the shape of the WAKE manifold (t-test on animal averages; mean point cloud radius: t(10) = 0.1, *P* > 0.05; point cloud standard deviation (SD): t(10) = 0.0, *P* > 0.05); **b**, HD cells in juvenile WT rats showed more improvement in HD tuning than those in *Fmr1^−/y^* rats when their tuning was estimated from the WAKE manifold rather than real HD (LME: genotype effect, P < 0.05). **c**, HD info of tuning curves estimated from the WAKE manifold was not dependent on the number of cells used to compute the manifold (r = 0.04, *P* > 0.05). **d,** Juvenile WT and *Fmr1^−/y^* rats had NREM epochs of similar duration (LME: no genotype effect, P > 0.05). **e**, Juvenile *Fmr1^−/y^* rats spent lower fraction of their sleep epochs in REM sleep than WT rats (t(10) = 2.59, *P* < 0.05). **f**, Mean firing rates of recorded cells during WAKE and REM are strongly correlated in both groups (WT: r = 0.84, *P* < 10^−87^; *Fmr1^−/y^*: r = 0.93, *P* < 10^−102^). **g,** There were no significant differences between groups in the shape of the REM manifold (t-test on animal averages; mean point cloud radius: t(10) = 0.81, *P* > 0.05; point cloud standard deviation (SD): t(10) = 1.30, *P* > 0.05); **h**, HD cells in juvenile WT rats showed more improvement in HD tuning than those in *Fmr1^−/y^* rats when their tuning was estimated from the REM manifold rather than real HD (LME: genotype effect, P < 0.01). **i**, HD info of tuning curves estimated from the REM manifold was not dependent on the number of cells used to compute the manifold (r = 0.19, *P* > 0.05). Dashed lines on histograms, population medians; horizontal lines on animal average panels, means of animal averages.

**Figure S4.**
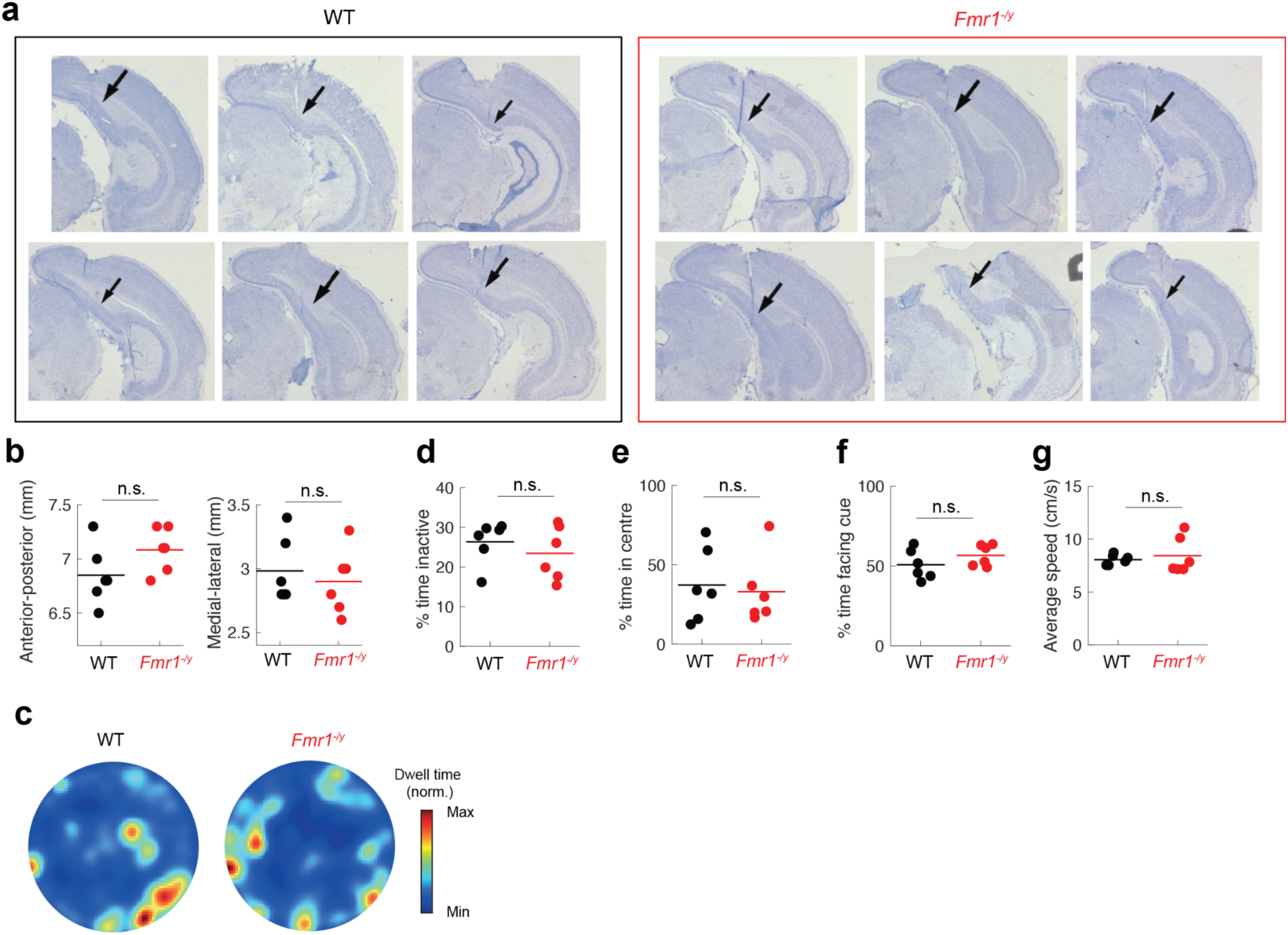
Probe placement and behaviour in adult WT and *Fmr1^−/y^* rats. **a**, Silicon probe tracts going through PoSub, as visualized by Nissl staining. Black arrows, electrode tracts. **b**, No significant differences in probe placement in the anterior-posterior (left) and medial-lateral (right) directions across genotypes (t-test on animal averages, anterior-posterior: t(10) = 1.67, *P* > 0.05; medial-lateral: t(10) = 0.58, *P* > 0.05). **c**, Heat maps of average occupancy across the two genotypes. **d**, Both groups spent equal amount of time inactive (t-test on animal averages: t(10) = 0.83, *P* > 0.05). **e**, Both groups spent the same amount of time in the centre of the arena (t-test on animal averages: t(10) = 0.33, *P* > 0.05). **f**, Adult *Fmr1^−/y^* rats spent the same amount of time as WT rats facing the visual landmark (t-test on animal averages: t(10) = 1.27, *P* > 0.05). **g**, Adult WT and *Fmr1^−/y^* rats ambulated with the same average speed within the active period (t-test on animal averages: t(10) = 0.51, *P* > 0.05). Horizontal lines on animal average panels, means of animal averages.

**Figure S5.**
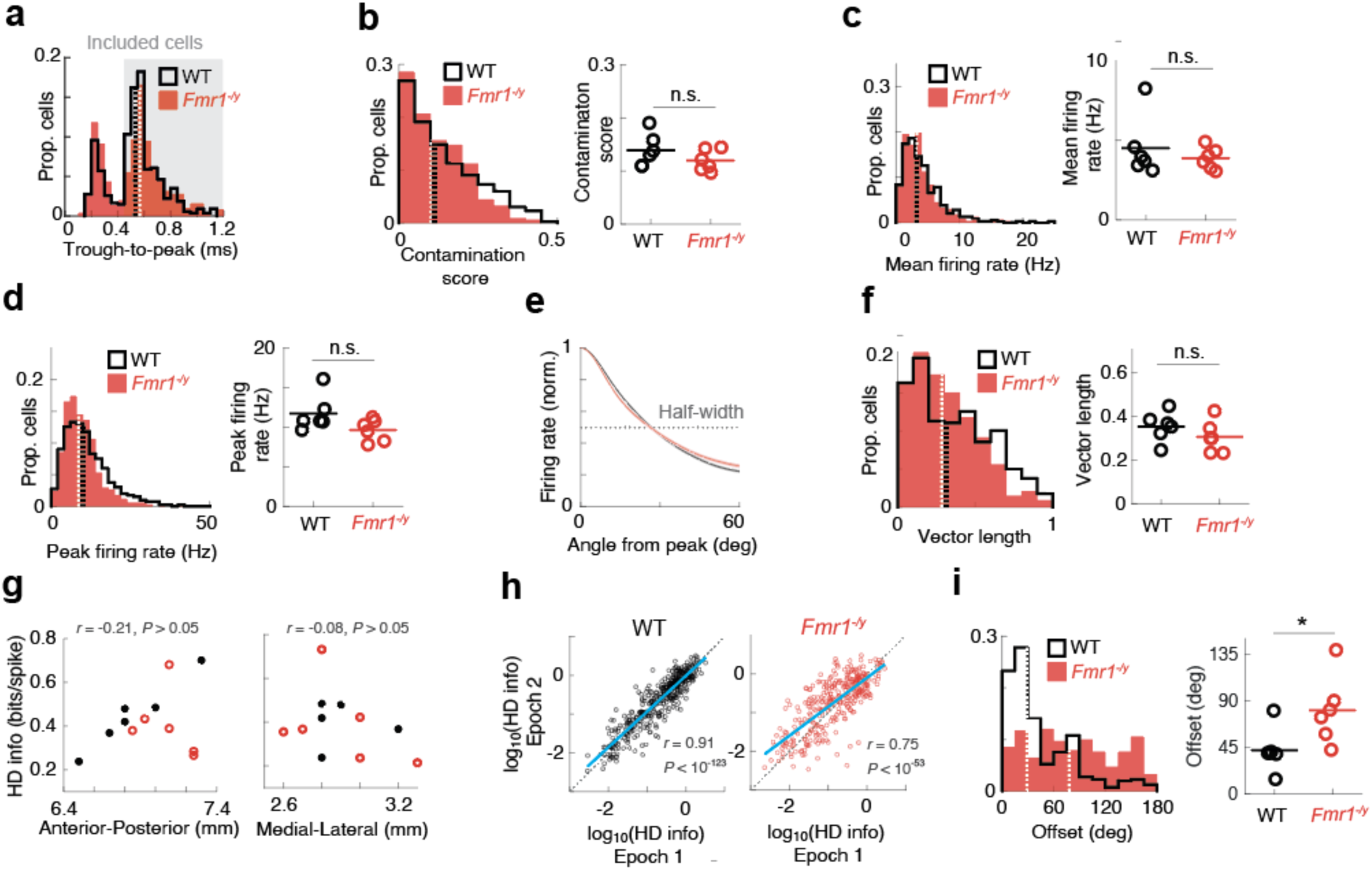
Single-cell and session metrics in adult WT and *Fmr1^−/y^* rats. **a**, Distribution of the through-to-peak duration of all recorded units. Grey background, cells classified as putative excitatory cells based on the trough in the bimodal distribution. **b**, Cluster contamination scores across the two groups (LME: no genotype effect, *P* > 0.05). **c**-**d**, Mean firing rates (LME: no genotype effect, *P* > 0.05) (**c**) and peak firing rates (**d**) of PoSub cells (LME: no genotype effect, *P* > 0.05). **e**, Mean normalized HD tuning curves of putative excitatory cells recorded in adult WT and *Fmr1^−/y^*rats. Only cells with tuning curve half-widths of less than 120 degrees were included. Shaded areas, +/- SEM. **f**, Resultant vector length across the genotypes (LME: no genotype effect, *P* > 0.05). **g**, No significant relationship between probe implantation coordinates and observed mean HD info for a given animal (anterior-posterior: r(10) = 0.21, P >0.05; medial-lateral: r(10) = 0.08, P > 0.05). **h**, HD tuning is preserved across the two exploration epochs in both genotypes (WT, *r* = 0.91, *P* < 10^−123^; *Fmr1^−/y^*, *r* = 0.75, *P* < 10^−53^). **i,** Adult *Fmr1^+/-^* rats display more drift in the anchoring of their HD tuning curves between two exploration epochs (LME: genotype effect, *P* < 0.05). Dashed lines on histograms, population medians; horizontal lines on animal average panels, means of animal averages.

**Figure S6.**
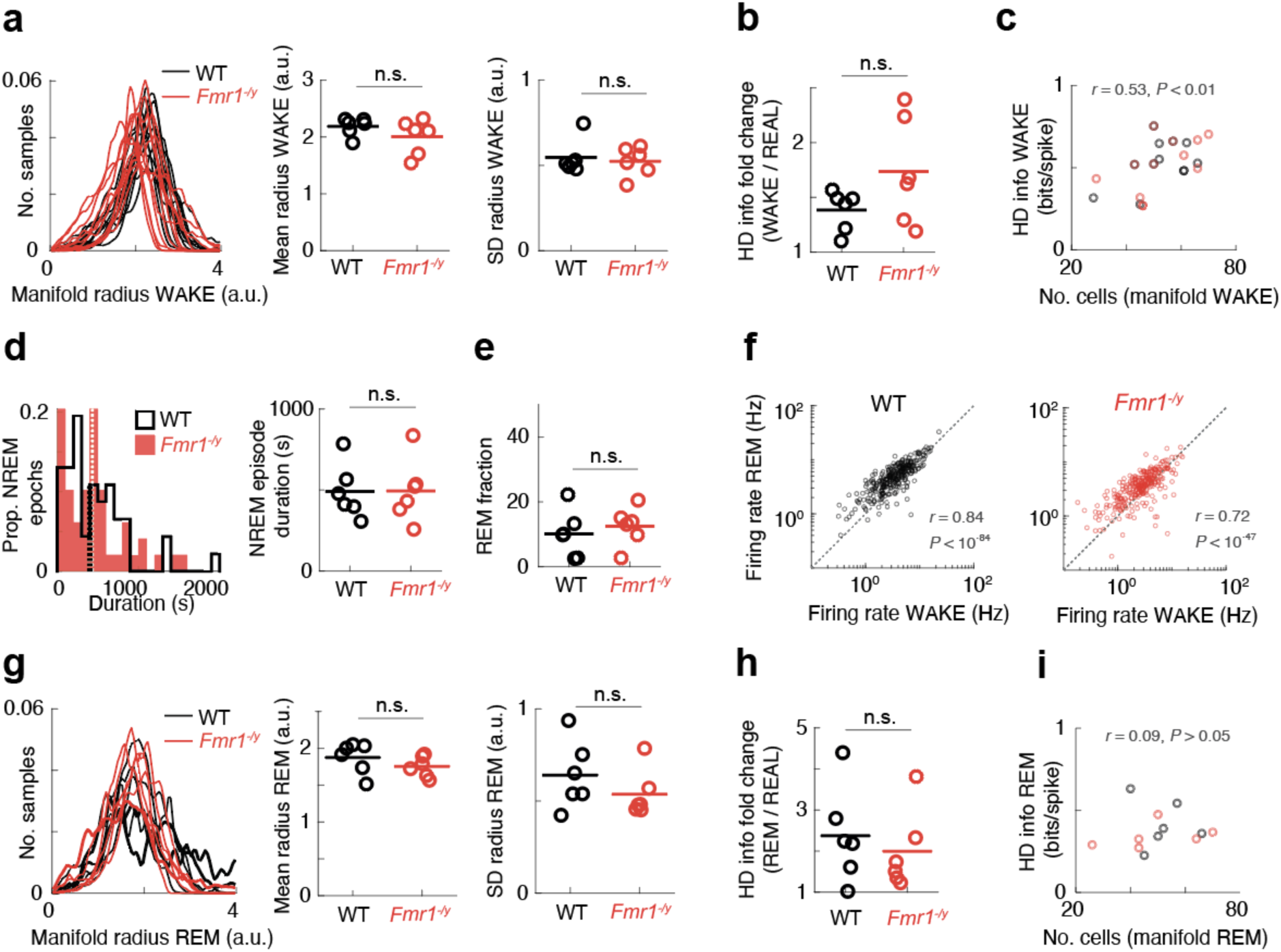
Manifold and sleep metrics in adult WT and *Fmr1^−/y^* rats. **a,** There were no significant differences between groups in the shape of the WAKE manifold (t-test on animal averages; mean point cloud radius: t(10) = 1.28, *P* > 0.05; point cloud standard deviation (SD): t(10) = 0.42, *P* > 0.05); **b**, PoSub cells in adult WT rats showed more improvement in HD tuning than those in adult *Fmr1^−/y^* rats when their tuning was estimated from the WAKE manifold rather than real HD (LME: genotype effect, P < 0.05). **c**, HD info of tuning curves estimated from the WAKE manifold was correlated with the number of cells used to compute the manifold (r = 0.53, *P* < 0.01). **d,** Adult WT and *Fmr1^−/y^* rats had NREM epochs of similar duration (LME: no genotype effect, P > 0.05). **e**, Adult *Fmr1^−/y^* rats spent the same fraction of sleep epochs in REM sleep as WT rats (t(10) = 0.63, *P* > 0.05). **f**, Mean firing rates of recorded cells during WAKE and REM were strongly correlated in both groups (WT: r = 0.84, *P* < 10^−84^; *Fmr1^−/y^*: r = 0.72, *P* < 10^−47^). **g,** There were no significant differences between groups in the shape of the REM manifold (t-test on animal averages; mean point cloud radius: t(10) = 1.18, *P* > 0.05; point cloud standard deviation (SD): t(10) = 1.13, *P* > 0.05); **h**, PoSub cells in adult WT rats showed the same amount of improvement in HD tuning as those in adult *Fmr1^−/y^* rats when their tuning was estimated from the REM manifold rather than real HD (LME: no genotype effect, P > 0.0). **i**, HD info of tuning curves estimated from the REM manifold was not dependent on the number of cells used to compute the manifold (r = 0.09, *P* > 0.05). Dashed lines on histograms, population medians; horizontal lines on animal average panels, means of animal averages. Dashed lines on histograms, population medians; horizontal lines on animal average panels, means of animal averages.

